# 20 years of African Neuroscience: Waking a sleeping giant

**DOI:** 10.1101/2020.06.03.131391

**Authors:** MB Maina, U Ahmad, HA Ibrahim, SK Hamidu, FE Nasr, AT Salihu, AI Abushouk, M Abdurrazak, MA Awadelkareem, A Amin, A Imam, ID Akinrinade, AH Yakubu, IA Azeez, GM Yunusa, AA Adamu, HB Ibrahim, AM Bukar, AU Yaro, LL Prieto-Godino, T Baden

**Author notes:** Correspondence: MBM, LLPG,TB.

## Abstract

Understanding the function and dysfunction of the brain remains one of the key challenges of our time. However, an overwhelming majority of brain research is carried out in the Global North, by a minority of well-funded and intimately interconnected labs. In contrast, with an estimated one neuroscientist per million people in Africa, news about neuroscience research from the Global South remains sparse. Clearly, devising new policies to boost Africa’s neuroscience landscape is imperative. However, the policy must be based on accurate data, which is largely lacking. Such data must reflect the extreme heterogeneity of research outputs across the continent’s 54 countries distributed over an area larger than USA, Europe and China combined. Here, we analysed all of Africa’s Neuroscience output over the past 21 years. Uniquely, we individually verified in each of 12,326 publications that the work was indeed performed in Africa and led by African-based researchers. This step is critical: previous estimates grossly inflated figures, because many of Africa’s high-visibility publications are in fact the result of internationally led collaborations, with most work done outside of Africa. The remaining number of African-led Neuroscience publications was 5,219, on average only ~5 per country and year. From here, we extracted metrics such as the journal and citations, as well as detailed information on funding, international collaborations and the techniques and model systems used. We link these metrics to demographic data and indicators of mobility and economy. For reference, we also extracted the same metrics from 220 randomly selected publications each from the UK, USA, Australia, Japan and Brazil. Our unique dataset allows us to gain accurate and in-depth information on the current state of African Neuroscience research, and to put it into a global context. This in turn allows us to make actionable recommendations on how African research might best be supported in the future.

## INTRODUCTION

Africa accounts for 15% of the global population but 25% of the global disease burden^1^. Moreover, the continent has the world’s largest human genetic diversity, with important implications for understanding human diseases^2^, including neurological disorders^3, 4^. However, even though early progress in neuroscience began in Egypt^5^, Africa’s research capacity remains weak^6^. The reasons for this are diverse, and include low funding^1^, inadequate research infrastructure ^7^, the relatively small number of active scientists^8^, and their overbearing administrative and teaching load^9, 10^. These barriers limit research and innovations from Africa^11^, and contribute to brain drain^12^.

Over recent decades, an increasing number of local and international initiatives were set-up to address some of these challenges, especially in neuroscience^13^. This seems to have led to some progress, as for example seen in a steady rise in the number of publications in the continent’s traditional science powerhouses such as South Africa, Egypt or Nigeria^14^. Of these publications, almost 70% have non-African based authors^6^. While on the one hand this may be indicative of important collaborative links between Africa and the rest of the world, it leaves it unclear which studies were truly African-led, and carried out in African labs – and which were rather led by researchers elsewhere^6, 15^. Indeed, a previous estimate suggested that as much as 80% of published health research with African authors was not African led^16^.

Database mining approaches using a combination of search terms such as ‘Neuroscience’ and ‘Africa’ have been used to estimate neuroscience research outputs from Africa^14, 17^. However, this approach does not delineate African-led studies from those led by researchers elsewhere. For example, PubMed data mining identifies 1,247 Nigerian neuroscience papers between 1996 and 2017. However, manual curation revealed that of those, 54% were led by non-Nigerian laboratories ^15^. The remaining 46% Nigerian-led studies generally had low visibility, with the majority being published in African journals that attract few non-African citations. This was complemented by a general absence of genetically modified model systems in these publications, and only occasional use of basic modern techniques such as fluorescence microscopy or western blotting^15^.

Clearly, these numbers are alarming. However, in view of the continent’s vast geographical, political and cultural diversity, extrapolating from Nigeria’s research output to the rest of the continent is not possible. Accordingly, to survey African neuroscience as a whole, we here manually went through each of 12,326 PubMed-listed African neuroscience publications since 1996. We again identified those that presented clear evidence that the research was indeed carried out in Africa (Methods), which eliminated ~58% of publications to leave 5,219 – on average only five per country and year. From here, we extracted key metrics including author affiliations, field of neuroscience, journals and citations, as well as information on funding, models, and techniques. For comparison, we also extracted the same metrics from 220 randomly selected publications each from the UK, USA, Australia, Japan and Brazil, of which 79% passed our inclusion criteria. We here present a summary of our main findings. All data is available at https://github.com/BadenLab/AfricanNeuroscience.

## RESULTS

### African neuroscience by the numbers

Africa’s Neuroscience output since 1997 has been dominated by a handful of countries: Egypt (n = 1,478, 28%), South Africa (n = 1,181, 23%), Nigeria (n = 566, 11%), Morocco (n = 409, 8%) and Tunisia (n = 388, 7%) (Fig. 1A). Together, these five scientific powerhouses account for more than three in every four neuroscience papers published from the continent. At 2-3% each, further contributions came from the East African nations of Kenya (n = 131), Ethiopia (n = 119) and Tanzania (n = 103), followed by 1-2% each from Cameroon (n = 81), Malawi (n = 71), Algeria and Senegal (n = 70 each), Uganda (n = 69) and Ghana (n = 60). Beyond these, numbers taper off further, with more than half of African countries contributing fewer than 10 papers. Nevertheless, over the past two decades, the number of Neuroscience publications published each year has exponentially increased across all of Africa’s major geopolitical regions (North, West, East, Central and Southern Africa, following the definition from the United Nations) (Fig. 1B). Accordingly, the continent’s neuroscience is on an all-time high, and one might expect this trend to continue going forward.

**Figure 1.**
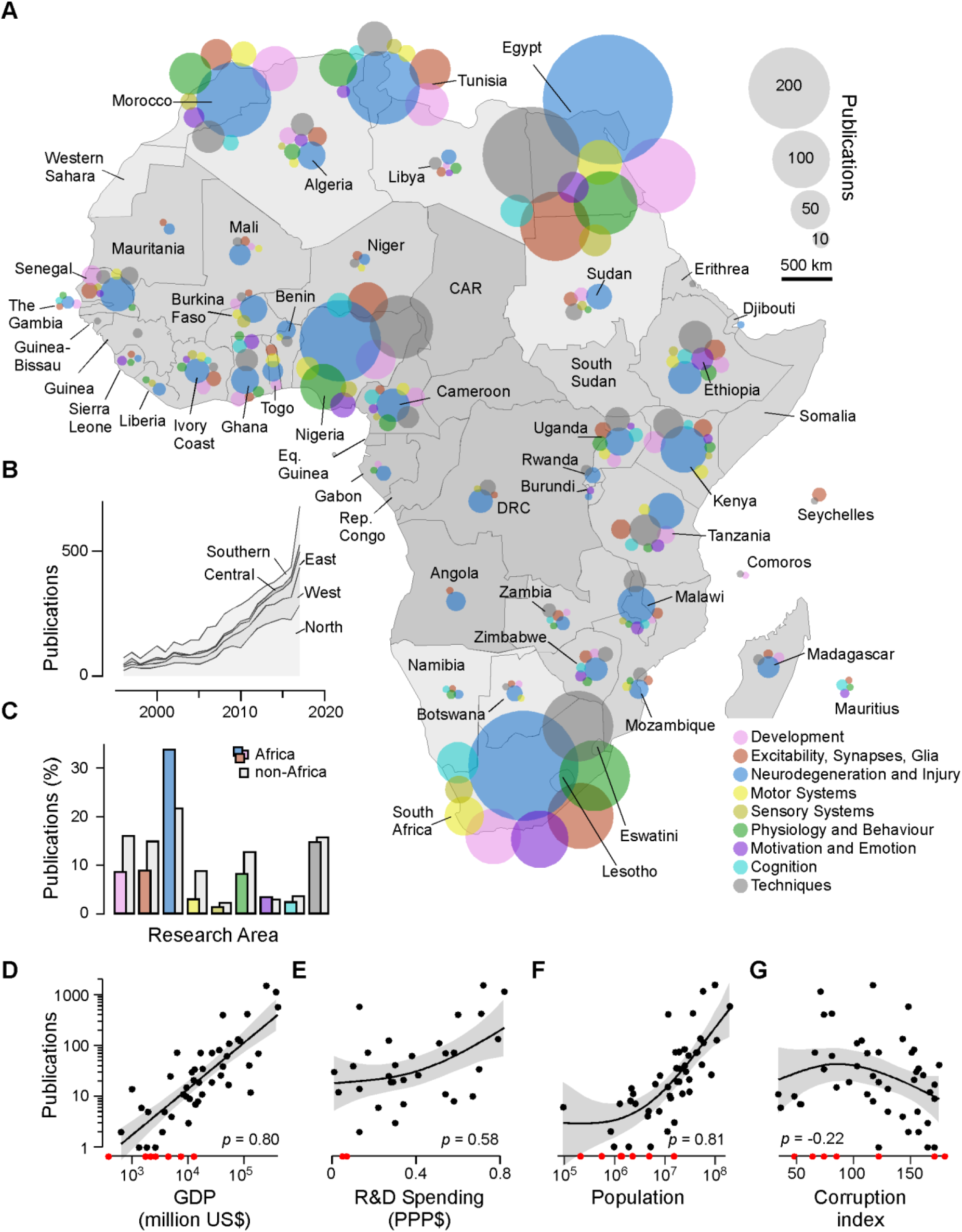
African Neuroscience 1996 – 2017. **A**, Overview of Africa’s neuroscience publications in the timeframe indicated, organised nine broad topics as indicated by the different colours. Bubble sizes denote the total number of papers per country and topic, as indicated. **B**, Total publications per year, with contributions from different African regions highlighted. Regions were delineated following the United Nations definition into North Africa, West Africa, East Africa, Central Africa and Southern Africa. See also background shading in (A). **C**, Distribution of research topics in Africa (coloured bars – for a legend see (A)) and outside of Africa (grey bars). **D-G**, Total number of publications per year and country plotted against gross domestic product (GDP, D), research and development (R&D) spending in purchasing power parity dollars (PPP$) (E), population (F) and corruption Index (G). Pearson-correleation coefficients as indicated. Data was fitted with generalized additive models (GAMs, solid lines, Methods), with shading indicating 95% confidence intervals. Data points in red indicate entries with n = 0 publications in the dataset.

Here, dominant research schemes^18^ include neurodegeneration and injury (n = 2,066, 34%; compared to 22% outside of Africa (OA)), followed by techniques (n = 905, 15%; OA: 16%), excitability, synapses and glia (n = 550, 9%, OA: 15%), development (n = 532, 9%; OA: 16%), and physiology and behaviour (n = 511, 8%; OA: 13%) (Fig. 1C). In comparison, research on motivation and emotion (n = 217, 4%; OA: 3%), motor systems (n = 191, 3%; OA: 9%), cognition (n = 155, 3%; OA: 4%) and sensory systems (n = 92, 2%; OA: 2%) is less prevalent. By and large, and despite a small degree of inevitable variation, this general distribution across major neuroscience research schemes has been surprisingly constant, both across countries (Fig. 1A), and over time (not shown).

To explore some of the basic factors that may contribute to a productive African Neuroscience research environment, we related each country’s total number of neuroscience publications to metrics of economy (GDP and R&D spending), population and governance (corruption index). This indicated that, perhaps unsurprisingly, a combination of money and manpower is a key requirement (Fig. 1D-F). A weak negative correlation to a country’s corruption index further supports this view (Fig. 1G, see also^19^).

### The visibility of African neuroscience

We next assessed the international visibility and communication of Africa’s neuroscience research outputs by way of citation numbers and journal impact factor (IF). For global context, we computed the same metrics for ~200 randomly selected papers each from the USA, UK, Japan, Australia and Brazil. This revealed a great diversity of African research visibility, with many papers ranking on par with many non-African papers (Fig. 2A). However, different regions varied markedly in the distribution of these metrics. For example, with a mean of ~13 citations per paper, West African publications tended to be cited least frequently. In contrast, Southern Africa’s publications were on average cited 31 times, on par with those coming from Brazil. Nevertheless, though dominated by the Global North (here: UK, USA, Japan, Australia, mean of ~77 citations per paper), also researchers from most African regions published at least a small fraction of papers in top bracket (citations ≥ 95), which presumably constitute our discipline’s most influential work. These trends were largely mirrored also in the publishing journal’s IF (Fig. 2B, C). However, unlike for citations, very few African publications ranked in the top bracket (here: IF ≥9.5), reiterating many of the frequently discussed shortcomings of this metric^20, 21^. Together, even though for now much of global neuroscience research remains dominated by the Global North, African neuroscience’s influence on the world’s production of knowledge is undeniably growing, with different regions representing different stepping stones along this journey.

**Figure 2.**
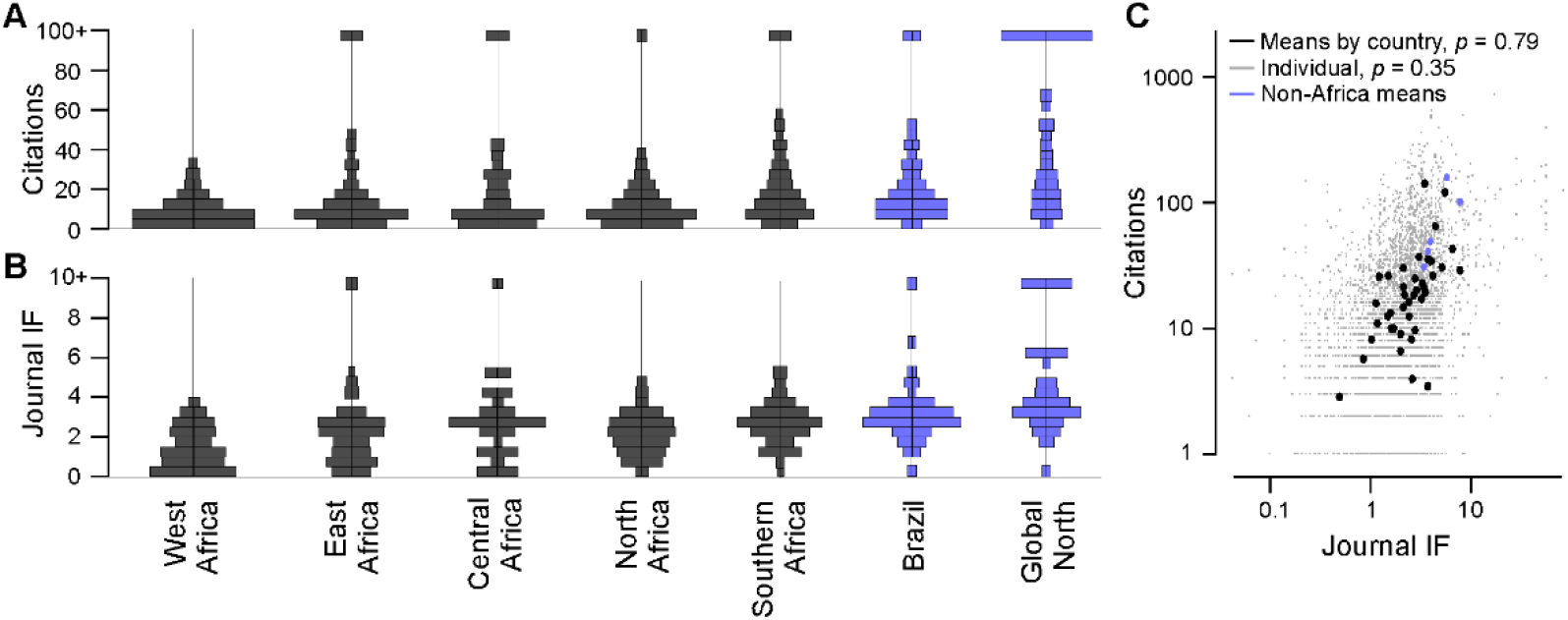
Citations and Impact. **A,B,** Area-normalised histograms of citations (A) and the publishing journal’s impact factor (IF, B) of all papers from different African and non-African regions, as indicated. In each case, all citations above ≥95, and IF above ≥9.5 were allocated to a single bin (top). **C**, Citations plotted against IF for every paper in the databased (small grey dots), and for means by country (large dots), as indicated. Linear correlation coefficients as indicated.

### International collaborations

One key aspect of integration into the global research community comes through international scientific collaborations. Here, the lack of funding and barriers related to visa-processes have long made it difficult for many African researchers to engage with colleagues abroad^22^. However, where these difficulties have been overcome, the visibility of research does stand to gain. For example, African-led neuroscience publications resultant from international collaborations – both within Africa and beyond – tended to be cited more frequently, and were published in higher IF journals (Fig. 3A). Perhaps unsurprisingly, the capacity for international collaboration is therefore paramount for research visibility^23^. We therefore next investigated how Africa’s collaborative networks are geographically organised.

**Figure 3.**
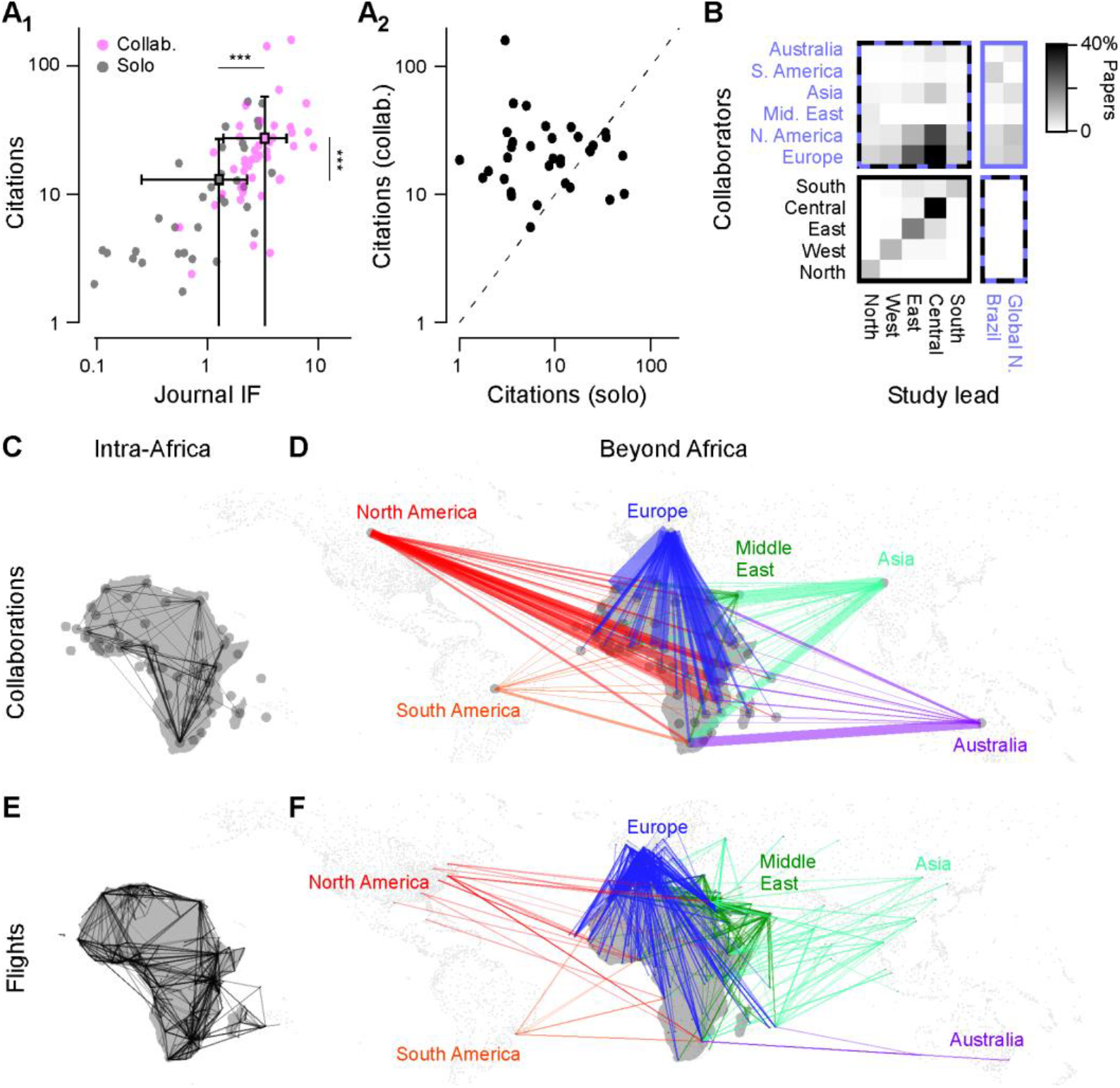
Collaborations. **A_1_**, Citations plotted against Journal impact factor (IF) for each African country, divided into publications without (grey, ‘solo’) and with international collaborations (pink, ‘collab.’). The latter included both within-African international collaborations as well as collaborations beyond Africa’s borders. Square-markers and errors denote each population’s mean and s.d.. Both citations and IFs were significantly higher for collaborative papers (Wilcoxon Rank Sum, 1 tailed, *** denoting p<0.001). **A_2_**, The same citation data as (A_1_) plotted pairwise for with (y-axis) and without collaborations (x-axis), highlighting the substantial positive influence of international collaborations. Dotted line indicates parity. **B**, Collaboration matrix between African regions and the rest of the world, with darker colours indicating a higher preponderance of collaborations. **C,D**, Intra-African (C) and Beyond-African (D) collaborative links organised by african country and major geopolitical regions beyond Africa, as indicated. The thickness of lines illustrates the total number of collaboratively authored papers, while colourings in (D) illustrate the collaboration partner beyond Africa. **E,F**, as (C,D), respectively, but for the existence of international flight routes based on data from OpenFlights.org. Each route is illustrated with a single line of consistent opacity and thickness.

This revealed that besides collaborating domestically, African Neuroscientists rarely collaborated across long distances *within* Africa – instead, the gaze was clearly pointed beyond the borders of the continent (Figs. 3B-D). In particular, many internationally co-authored papers had links with either Europe or North America. Similarly, any intra-Africa collaborations were mostly “looking upwards”. For example, West, East and Central African-led studies occasionally included co-authors from Southern Africa, while vice versa Southern African-led papers almost never had East- West- or Central African co-authors (Fig. 3B). All the while, North Africa kept mostly to itself with regards to pan-African collaborations, but instead mostly collaborated with Europeans, North Americans, and – uniquely for Africa – the Middle East. The latter collaborations follow a geographic and cultural pattern^24, 25^.

Overall, the striking preponderance of collaborations beyond Africa’s borders over pan-African international links is highly reminiscent of the continent’s logistical networks – here exemplified by available international flights (Fig. 3E,F). It seems likely that both networks are linked to common underlying factors such as historic, linguistic and cultural ties as well as economic considerations^25^. Nevetheless, this currently poor international connectedness within the continent ought to be considered in future efforts aiming to build a more united African research landscape.

### Funding African neuroscience

Who, if anyone, is funding Africa’s Neuroscience research? To investigate, we assessed funding declarations in Africa’s 265 top papers (Methods). Of these, many (n = 93, 35%) declared no funding at all (Fig. 4A). This lack of declarations was pervasive throughout the continent, but particularly prominent in Northern Africa (n = 46 of 73, 63%). It is likely that many of these studies were self-funded.

**Figure 4.**
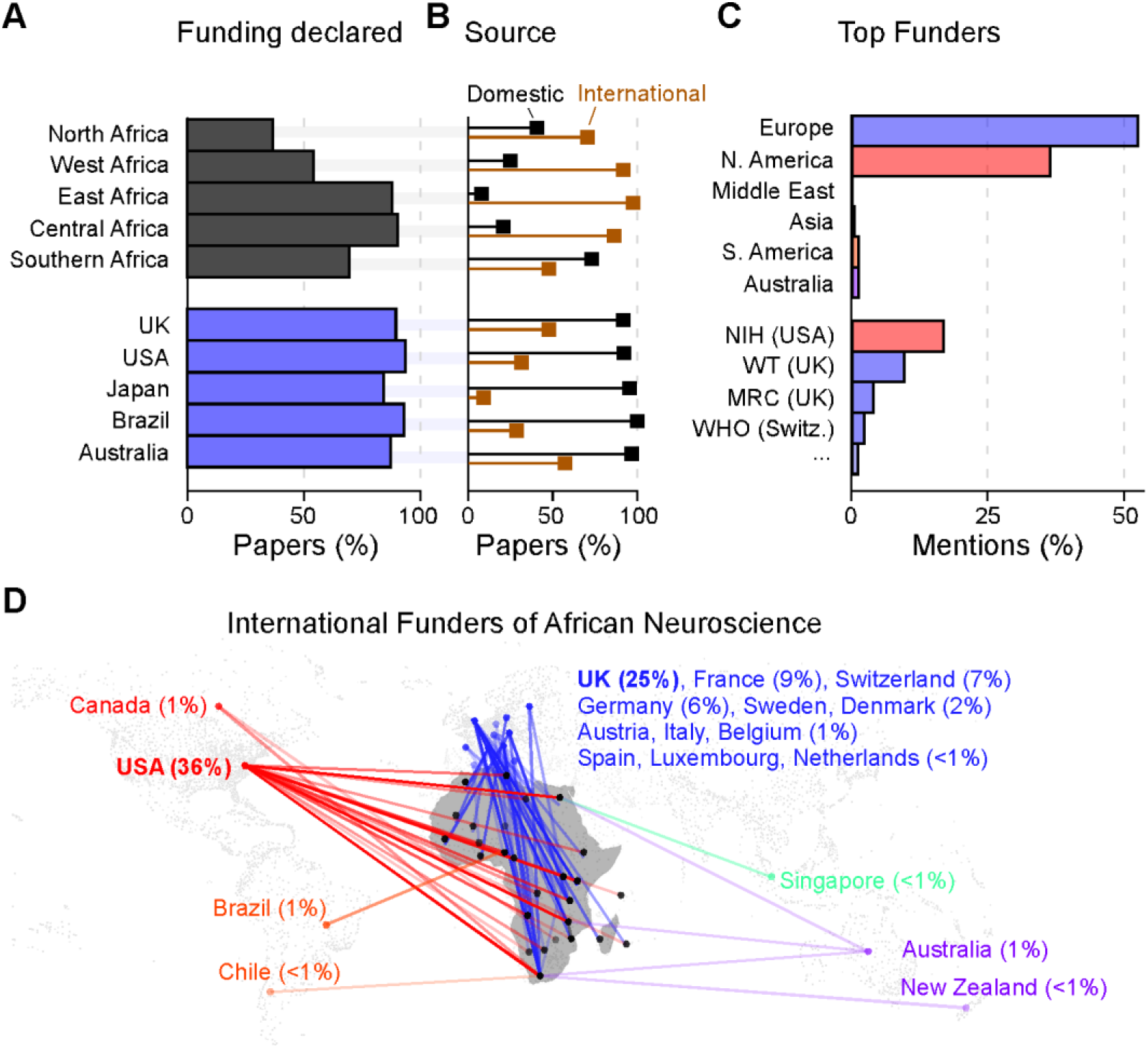
Funding. **A**, Percentage of papers with IF≥5 that included any form of funding acknowledgements. **B**, From (A), where declared, the source of funding, classed as domestic (black) and international (brown). International funding includes those received from any other country, including other African countries. Percentages are computed from each paper declaring either domestic or international support, or both. **C**, Percentages of funder mentions of all African papers with IF≥5 where international funding was declared (corresponding to the sum of brown columns in (B)). Where present, multiple funder mentions per single paper were individually included. **D**, International funding links from (B,C) displayed by geography. Each funding mention is illustrated with a single line of consistent opacity and thickness.

Of papers that did declare the funding (n = 172), we next assessed whether the sources were domestic and/or international (Fig. 4B). This revealed that most African neuroscience was supported by international rather than domestic agencies. For example, only 3 out of 37 (8%) of East African top papers declared domestic funding, while 36 (97%) declared international funding. The only African region where the number of domestic funding mentions exceeded international funder mentions was Southern Africa (n = 49 (73%) domestic, n = 32 (48%) international; of n = 67 total). In comparison, between 92% (UK) and 100% (Brazil) of papers included in our analysis declared domestic funding, with between 9% (Japan) and 57% (Australia) of papers declaring additional international support. It seems clear that the availability of local, rather than (or in addition to) international funding is critical to building a viable research culture, and Southern Africa appears to be the only region that is beginning to reflect this need. Indeed, South-Africa, by far Southern African’s largest research contributor, is the only country in Africa that invests nearly 1%, of its GDP in research and development, as recommended by the African Union in 2007^1^.

Nevertheless, with 46% (n = 123) of Africa’s 265 top neuroscience papers declaring international funding, external money being invested into Africa’s research landscape clearly did have important impact. The vast majority of this support came from the USA, who supported n = 44 (36%) of these 123 papers, followed first by the UK (n = 31, 25%), and then France (n = 11, 9%), Switzerland (n = 9, 7%) and Germany (n = 7, 6%) (Fig. 4C,D). Accordingly, unlike international collaborations (Fig. 3D), international funding support from the Middle East, Asia, Australia and South America for African neuroscience was essentially nonexistant. By agency, the USA’s NIH was acknowledged most frequently (n = 42, 34%), followed by the UK’s Wellcome Trust (n = 24, 20%) and Medical Research Council (n = 10, 8%). Already in a distant fourth place was the World Health Organisation (n = 6, 5%), and beyond this, no agency received more than 2% of international funding mentions.

### Model systems, techniques and medicinal plants

Racing advances in our understanding of nervous systems are notably driven by equally rapid advances in (bio)technology. Accordingly, access to state of the art research tools – both technological and biological – remain central to scientific success. Accordingly, understanding the availability and use of such tools across Africa is likely to be pivotal to any strategy to support future research. To this end, we categorised methods employed in each of the surveyed >6,000 papers as either “basic” or “advanced”. Specifically, if studies included any form of fluorescence microscopy, molecular biology or cell culture work, they were categorised as “advanced”. In contrast, studies based on classical histology, chromatography and/or behaviour were classed as “basic” (for a full list of inclusion and exclusion criteria, see methods). Despite this set of arguably conservative criteria, and with the notable exception of The Gambia (n = 5/14; 36%, all linked to an MRC-funded research unit), no African country’s neuroscience publications comprised more than a quarter of “advanced” entries (Fig. 5A_1,2_). Leading here were Egypt (n = 363/1,478; 25%), followed by South Africa (n=272/1,181; 23%), Morocco (n = 68/409; 17%), Tunisia (38/388; 10%); Nigeria (45/566; 8%), Ethiopia (9/119; 8%), and Algeria, (2/35; 6%). All other African countries ranked below 5% including many at 0% (countries with fewer than 10 papers were excluded from this analysis). In contrast, Japan, UK and USA all published 75% of papers based on “advanced” techniques, followed by Australia (54%) and Brazil (33%).

**Figure 5.**
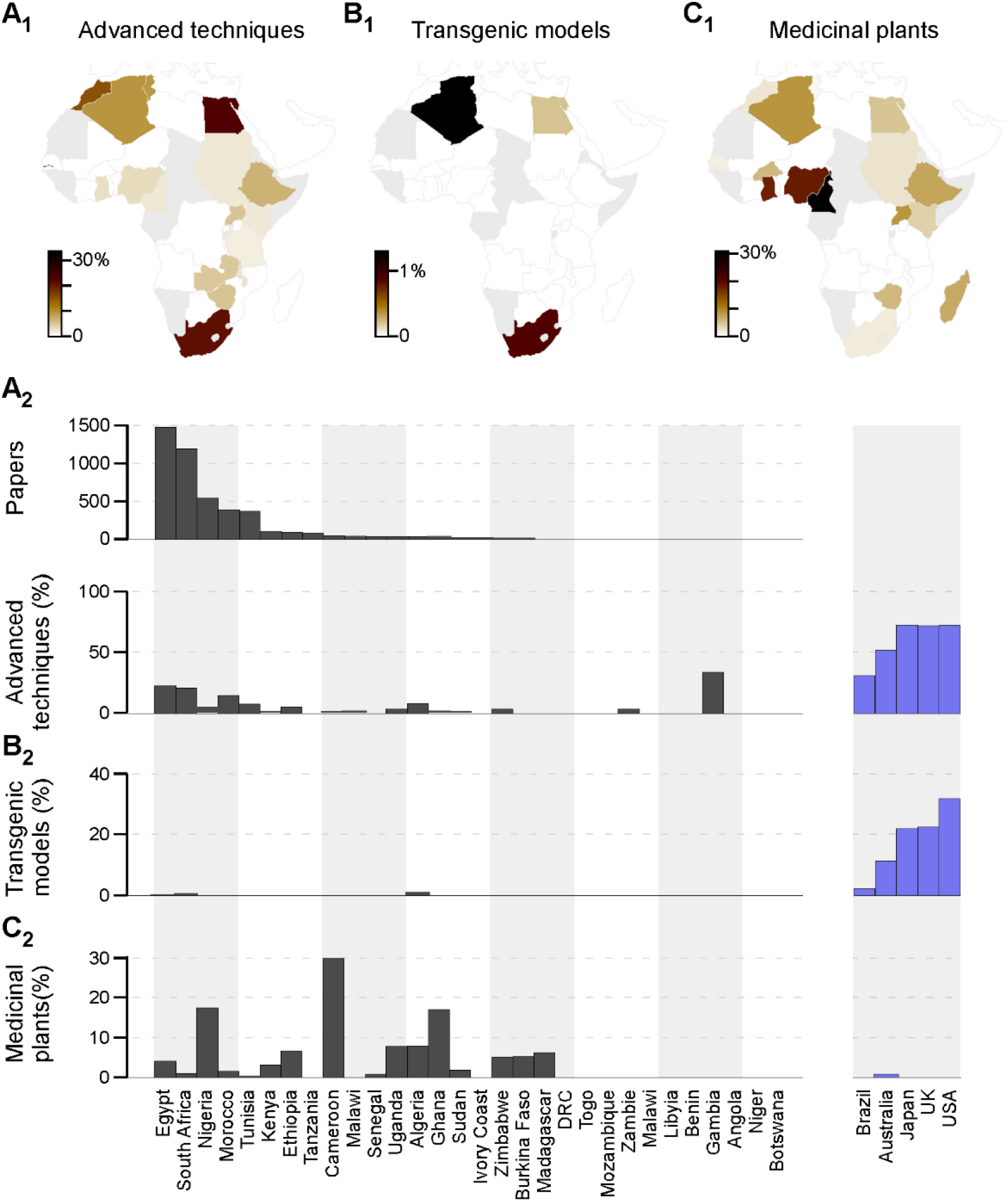
Research Techniques. **A_1_-C_1_,** Percentages of papers that used “advanced” techniques (Methods) (A_1_), transgenic models (B1) or medicinal plants (C_1_), organised by geography. Countries with fewer than 10 papers in the dataset were excluded (grey). **A_2_-C_2_**, As (A_1_-C_1_), plotted as percentage bars per country, with the same metrics extracted from representative non-African publications (blue). African countries are sorted by the total number of papers published (A_2_, top), excluding countries with fewer than 10 papers in the database.

Next, there was a near complete absence of small, low cost and genetically tractable model systems such as fruit flies, zebrafish or *C. elegans*^26^ in African neuroscience (Fig. 5B_1,2_). Unlike USA (33%), UK and Japan (23%), Australia (12%) or Brazil (3%), no African country used any genetically modified model systems (including cell culture or mice) in more than 1% of neuroscience publications. Most countries used none at all. Clearly, the promotion of the use of such model systems should be considered as part of strategies aimed to modernise Africa’s research landscape.

Finally, we assessed the use of endemic medicinal plants in African neuroscience research, many of which have been used for centuries for the treatment of diseases. Research in natural medicinal products puts Africa in an excellent position in the area of drug discovery ^27^. This revealed that research in this field is highly diverse across the continent (Fig. 5C_1,2_). In particular, several West-African countries with tropical and subtropical climates have invested heavily into this branch of Neuroscience, most notably Cameroon (30%) as well as Nigeria and Ghana (18% each). In contrast, many other countries, including Africa’s leading science powerhouses of Egypt (5%) and South Africa (1%), are more focussed on other topics.

## DISCUSSION

Our unique dataset highlights that Africa’s neuroscience productivity is on an all-time high, with a clear and ongoing upwards trajectory. Similarly, while the number of neuroscientists on the continent remains tiny compared to total population^28, 29, 30^, also the neuroscience scientific workforce is on the rise. This is for example mirrored in the increasing number of neuroscientists attending the Society of Neuroscientists of Africa (SONA) bi-annual meetings^31^, or a continuous rise in the number of applications for African-based neuroscience training programmes. However, to continue supporting this growth, major and ongoing investment into African science must be ensured.

Most declared neuroscience funding came from external sources, most notably from the USA and the UK. However, local funding is instead needed for establishing a viable African neuroscience research environment. At the moment, none of Africa’s 54 countries invests as much as 1% of their GDP into R&D, as recommended by the African Union^1^. South-Africa is one of the few countries nearly meeting this target, which likely part-explains its leading role in African neuroscience. For African research to continue to grow exponentially, African governments must step up and provide reliable support to their domestic research sector. Such efforts might also be well-supported by the local philanthropic sector. Africa has a large number of individuals and charitable organisations with access to substantial funds^32^. Governments, scientists and the general population must engage with these to contribute to local science funding, much like major non-African philanthropic organisations such as the Gates Foundation or Wellcome Trust that currently fund research on the African continent. To attract more funding, there is a need for African Neuroscientists to engage in Neuroscience advocacy campaigns to raise the profile of their research and its relevance, especially to local problems.

Although there is clear evidence of increasing Neuroscience outputs from African laboratories, Africa has much to do before it can catch up with the Global North. Based on citation and IF metrics, there remains substantial heterogeneity in the visibility of the neuroscience outputs across the continent. Under the caveat of using such metrics^20, 21^, West Africa seems to lag behind among all the regions. For example, the region’s giant – Nigeria, published only one Neuroscience paper in a ‘top‐tier’ journal in the 21-year period^15^. The lack of visibility, especially in citations, may be part-explained by choices over where work is submitted for publication. Many Nigerian Neuroscience papers are published in African journals, which generally offer poor visibility beyond the continent’s borders^15^. Moreover, many African journals are not PubMed indexed (and therefore excluded from our study)^33^. Given the clear benefit of publishing in indexed journals for driving research visibility and collaboration, this flags the need for African academics to increasingly target indexed journals. This will be facilitated by increasing the widespread availability of fee-waivers from international outlets^34^.

Next, our analysis highlights a profound lack of state-of-the-art equipment and modern experimental approaches. With few notable exceptions, basic tools like fluorescence microscopy, molecular biology or cell culture were used in less than 10% of most African countries’ Neuroscience publications (see also Ref^15^). For example, to our knowledge most African countries have no functional confocal microscope. Where present, with a handful of exceptions, equipment of this type tends to be associated with internationally well-connected (and typically funded) private research centres. Next, while some public institutions do have some “advanced” equipment located in individual departments, a frequent lack of willingness to share, even locally, can limit their wide-spread use^15, 34^. Although funding schemes and training programmes have enabled many African scientists to acquire modern neuroscience skills in foreign labs, the absence of the same research infrastructure back at their home institutions continues to restrict the extent to which such skills can be put into use. Clearly, beyond financial investment, African researchers must be afforded widespread access to modern research infrastructure^9^. In addition to the provision of training opportunities abroad, local and international Neuroscience funding initiatives should support African scientists to establish their laboratories. Similarly, African labs have much to gain from investing in infrastructure and expertise in designing and producing research-grade open hardware equipment^9, 35, 36, 37^.

Finally, the near-complete absence in the use of transgenic models in African Neuroscience is worrying, and likely contributes substantially to the generally low visibility of Africa’s neuroscience community. Instead, many African neuroscientists continue to rely on wild-type rodents models, most notably rats, followed by mice ^15^. The cheap and genetically amenable nature of model systems like zebrafish, fruit flies or *C. elegans*, makes these models ideal for African Neuroscience compared to many mammalian models. One-third of the Nobel Prizes in Physiology and Medicine awarded between 1996 and 2017 relied heavily on non‐mammalian yet genetically accessible model systems^38^. The many challenges faced by African Neuroscience, most notably lack of funding, make ultra-low-cost models like fruit flies and *C. elegans* particularly interesting for research on the continent^39^. This particularly calls for increased investment to facilitate the use of these and other similar affordable and genetically amenable model species in African neuroscience. For this, scientists and funding agencies will also need to work closely with national goverments and biosafety authorities to put regulation for the import and use of genetically modifiable animals in place, which to date is missing in many African countries.

## CONCLUSIONS

Taken together, while African Neuroscience remains comparatively small, it is clearly on the rise. To sustain this rise and increase the continent’s neuroscience visibility, there is a clear need for increased investment in modern research equipment, training in the use of this equipment, and the adoption of genetically tractable models. While some of this investment will likely continue to come from beyond Africa’s borders, it will be critical to bolster African countries’ domestic research support streams, from governments and private funders alike. Next, while international collaborations are valuable, African neuroscience must in parallel be strengthened through intra-African collaborations and the promotion of sharing of restricted resources.

In view of the highly interdisciplinary nature of Neuroscience, many aspects of our findings are likely to generalise to other scientific disciplines, ultimately highlighting important implications for future milestones in Africa’s path to becoming a veritable giant in the world’s generation of scientific knowledge.

## Acknowledgments

We thank Philipp Berens for computing GAMs for Figure 1. Funding was provided by the European Research Council (ERC-StG “NeuroVisEco” 677687 to TB, ERC-StG “EvolutioNeuroCircuit” 802531 to LLPG), The UKRI (BBSRC, BB/R014817/1 to TB, and MRC, MC_PC_15071 to TB), the Leverhulme Trust (PLP-2017-005 to TB), the Lister Institute for Preventive Medicine (to TB). LLPG’s research was supported by The Francis Crick Institute. We also wish to thank the FENS-KAVLI Network of Excellence for support (TB, LLPG), as well the as EMBO YIP programme (TB).

## Author Contributions

The study was conceived and organised by MBM. Data curation was done by MBM, AU, IHA, HSK, NFE, SAT, AAI, AM, AMA, AA, IA, IA, AID, YAH, AIA, YGM, AAA, IHB, BAM, and YAU and managed by MBM.

Data analysis was done by TB. The paper was written by MBM, LLPG and TB.

## METHODS

### Data availability

All data used in the study is freely available on GitHub: https://github.com/BadenLab/AfricanNeuroscience.

### Data extraction

Neuroscience-related research articles from Africa, USA, UK, Australia, Japan or Brazil from January 1996 to December 2017 were retrieved from PubMed. Search terms used were “Neuroscience” OR “Nervous system” OR “Brain” OR “Neuron” OR "Spinal cord", in combination with the name of each of the individual African and non-African countries were used. Of these, primary research, case reports or clinical trials were included, while review articles were excluded. Next, duplicates and irrelevant articles were manually removed. This yielded a total of 12,326 candidate papers from Africa. For comparison, 220 papers each from the abovelisted non-African countries were also analysed, after randomly selecting 10 publications per year and country using the same search terms. Of these total of 1,100, n = 229 (21%) were eliminated based on the same exclusion criteria applied to our African dataset to leave a total of n = 871 non-African papers (Australia: 164; Brazil: 173; Japan: 197; UK: 171; USA: 166)

### Data curation

To identify research conducted within each country, the full texts of all the articles were retrieved and screened manually. For exclusion, papers from outside of Africa were identified based on the listed affiliations of lead/corresponding/senior author(s) as well as study location. The latter was extracted from information in the materials and methods or acknowledgements, where possible. For example, articles with external collaborations in which only a small fraction of the work was conducted within Africa, such as sample collection, were excluded. This process eliminated n = 7,107 papers, leaving n = 5,219 African papers for further analysis.

The latter were further screened by hand to retrieve the total number of google scholar citations, the publishing journal and its Clarivate Analytics impact factor, as well as information on model species whether or not they used medicinal plants. Impact factor for journals not indexed by Clarivate Analytics were estimated from Scimago. In addition, author affiliations were screened for evidence of collaboration (between research institutes, both nationally and internationally). In addition, we summarised each paper’s research techniques as either “basic” or “advanced”: Advanced techniques included electron microscopy, western blotting, immunohistochemistry, cell culture techniques, cloning, flow cytometry, fluorescence microscopy, whole-brain imaging, sequencing and identifying genes of interest, molecular cloning and recombinant DNA technology, gene delivery strategies, making and using transgenic organisms, manipulating endogenous genes, as well as any additional technique that was judged to be similarly advanced, where required. Basic techniques included histology, biochemical assays, such as enzyme‐linked immunosorbent assay (ELISA), plant extract preparation, high‐performance liquid chromatography (HPLC), behaviour and blood analysis. A paper was classed as advanced if it used any advanced technique, even if it mainly used basic techniques. In addition, each paper was attributed to one of nine broad topical themes, as put forward by the Society of Neuroscience (see also ^18^. Specifically, topics included (i) techniques, (ii) cognition, (iii) motivation and emotion, (iv) physiology and behaviour, (v) motor systems, (vi) sensory systems, (vii) neurodegeneration and injury, (viii) excitability, synapses and glia, and (ix) development. Finally, for all n = 265 African and n = 232 non-African papers that were published in journals with an impact factor ≥5, we also extracted information on funding. Data on international flights was taken from OpenFlights.org. Data and sources for other metrics, including GDP, R&D spending, GERD, population etc are detailed in the raw data tables provided on GitHub (see data availability). Starting from raw Microsoft Excel tables provided, all data analysis was performed in Igor Pro 6 (Wavemetrics) and GNU-R.

### Generalized additive models (GAMS)

For Fig. 1D-G, smooths were fit to the scatter plots using the lm-method (GDP) or the gam-method with the R-package ggplot. GAMs are generalized additive models, allowing to model nonlinear relationships between the variable on the x-axis and the variable of the y-axis modeled through spline basis functions. Grey areas correspond to 95%-CIs. For the GDP, performing model selection using the R-package mgcv indicated that the data did not support a nonlinear relationship.

## Notes

https://github.com/BadenLab/AfricanNeuroscience

### Competing Interest Statement

The authors have declared no competing interest.

